# MAP: a comprehensive pipeline for mobilome annotation and cargo gene characterisation in prokaryotic (meta)genomic assemblies

**DOI:** 10.64898/2026.09.11.750868

**Authors:** Alejandra Escobar-Zepeda, Martin Beracochea, Tatiana A. Gurbich, Paul Wilmes, Robert D. Finn

**Affiliations:** EMBL-EBI, Microbiome Informatics Team, Wellcome Genome Campus, Hinxton, Cambridge CB10 1SA, United Kingdom; Luxembourg Centre for Systems Biomedicine, University of Luxembourg, L-4362 Esch-sur-Alzette, Luxembourg; Department of Health, Medicine and Life Sciences, Faculty of Science, Technology and Medicine, University of Luxembourg, L-4362 Esch-sur-Alzette, Luxembourg

## Abstract

**Summary:** Mobile genetic elements (MGEs) drive horizontal gene transfer in prokaryotes, disseminating antimicrobial resistance genes (ARGs), virulence factors (VFs) and biosynthetic gene clusters (BGCs). Given their importance, there is a pressing need for a single, open source tool that annotates the MGE repertoire together with its functional cargo. We present MAP (Mobilome Annotation Pipeline), a Nextflow pipeline that predicts plasmids, viral sequences, prophages, integrons, insertion sequences, transposons, integrative and conjugative elements, and non-autonomous compositional outliers, removes redundant predictions, and labels genes within MGE boundaries. MAP outputs a GFF3 formatted file, a FASTA file of MGE sequences, and a combined report placing ARGs, VFs, toxins and BGCs in their mobilome context, enabling the identification of composite elements such as ARG-carrying integrons within plasmids. We demonstrate its use on genomes from the MGnify soil genome catalogue.

**Availability and Implementation:** MAP is written in Nextflow and Python under the Apache 2.0 licence and is freely available at https://github.com/EBI-Metagenomics/mobilome-annotation-pipeline and https://workflowhub.eu/workflows/452

## 1 Introduction

Mobile genetic elements (MGEs) are major drivers of horizontal gene transfer in prokaryotes, mediating the exchange of adaptive traits including antimicrobial resistance, virulence and biosynthetic capabilities across microbial communities (Carr et al. 2021; Haudiquet et al. 2022). They span a broad diversity of forms — plasmids, phages and prophages, integrons, insertion sequences, transposons, and other mobilisable elements — some autonomous, others non-autonomous, mobilised in trans by enzymes encoded elsewhere and recognisable only by their compositional bias. MGEs frequently recombine into composite elements, such as phage-plasmids or insertion sequences in integrons nested within plasmids (Pfeifer et al. 2022). Accurately annotating these composite elements requires resolving multiple element types and their genomic context simultaneously.

Although many tools predict individual MGE classes, no comprehensive open-source solution performs *de novo* annotation of a broad range of MGE repertoires together with their functional cargo across diverse datasets. Existing tools each tackle part of this task: proMGE detects MGEs from recombinase profiles (Khedkar et al. 2022), MobileElementFinder matches a curated database of known elements (Johansson et al. 2021), MOBHunter predicts elements from composition and marker genes (Rojas-Villalobos et al. 2025), and DeepMobilome screens read alignments against user-chosen target sequences (Cho et al. 2025). Most of these tools run only as web servers, limiting high-throughput analysis, and each covers a subset of MGE types with little functional annotation of cargos. To close this gap, we developed the Mobilome Annotation Pipeline (MAP) providing *de novo*, command-line annotation of diverse autonomous and non-autonomous MGE types, including plasmids, phages, integrons, insertion sequences, transposons, integrative and conjugative/mobilisable elements, and compositional outliers. MAP filters low-quality predictions, resolves redundancy, and annotates cargo genes within MGE boundaries. The tool additionally runs functional-annotation subworkflows for virulence and toxins, antimicrobial resistance, and biosynthetic gene clusters (BGCs), producing an integrated report using a format similar to that of PathoFact2 (Delgado et al. 2026), contextualising these cargo genes within the mobilome and BGCs. The MAP is part of the MGnify resource (Richardson et al. 2023).

## 2 Pipeline description

### 2.1 Installation and dependencies

MAP requires only Nextflow and a container engine (e.g. Singularity/Apptainer or Docker). Reference databases are fetched automatically (--download_dbs) or supplied through a configuration file. The pipeline follows nf-core best practices (Ewels et al. 2020) and FAIR principles (Wilkinson et al. 2016), is fully containerised and versioned, and records all tool and database versions at runtime.

### 2.2 Input files

MAP takes a CSV file listing a sample identifier and FASTA file path for each assembly. Optionally, users may provide pre-computed proteins (paired GFF and FASTA files), VIRify v4.0.0 or later (Rangel-Pineros et al. 2023) predictions, and an InterProScan (Blum et al. 2026) output table, allowing MAP to skip redundant computation when the assemblies were previously annotated, for example by mettanotator (Gurbich et al. 2025).

### 2.3 Annotation workflow

Contigs are renamed and filtered by length according to the requirements of each downstream tool; genes are predicted with Prodigal (Hyatt et al. 2010) and tRNAs with ARAGORN (Laslett and Canback 2004) as shown in Figure 1 section 1. MGEs are then predicted in parallel: geNomad (Camargo et al. 2024) detects plasmids, viruses and prophages (predictions retained above geNomad score of 0.8); ISEScan (Xie and Tang 2017) detects insertion sequences (only predictions flanked by inverted repeats are retained); IntegronFinder (Néron et al. 2022) detects integrons (only integrons labelled as complete are retained); and an ICEfinder2-lite subworkflow —refactored from ICEfinder2 (Wang et al. 2024) — annotates integrative and conjugative elements (ICEs) and integrative and mobilisable elements (IMEs). A compositional-outlier module detects non-autonomous elements on contigs of length >=100 kb using GC and k-mer-based z-scores, then refines boundaries using flanking repeats (Figure 1, section 2). In addition, geNomad predictions can be complemented with VIRify results: where the two tool outputs overlap by >25% of the shorter sequence (otherwise, the two independent predictions on the same contig are retained), geNomad prediction takes precedence unless a ViPhOG (Moreno-Gallego and Reyes 2021) annotation is present, and contigs classified as plasmid by geNomad and phage by VIRify are labelled as phage-plasmids. During integration, mobile genetic elements shorter than 500 bp or lacking coding sequences in autonomous MGEs are discarded as we expect to detect at a minimum the mobilisation enzymes. Compositional outliers overlapping tRNAs are removed and to avoid redundant entries in the GFF file, compositional outliers overlapping >= 75% to other MGEs are discarded, with all exclusions logged. CDSs overlapping an MGE by >90% of their length are labelled mobilome-resident and the original contig names are restored (steps depicted on section 3 of Figure 1). Final results are written in GFF3 format, which is validated using the GFF3 validator in GenomeTools (Gremme et al. 2013). In addition, a FASTA file of the MGE sequences is generated. Three derived GFFs additionally report passenger CDSs and their functional annotations.

**Figure 1.**
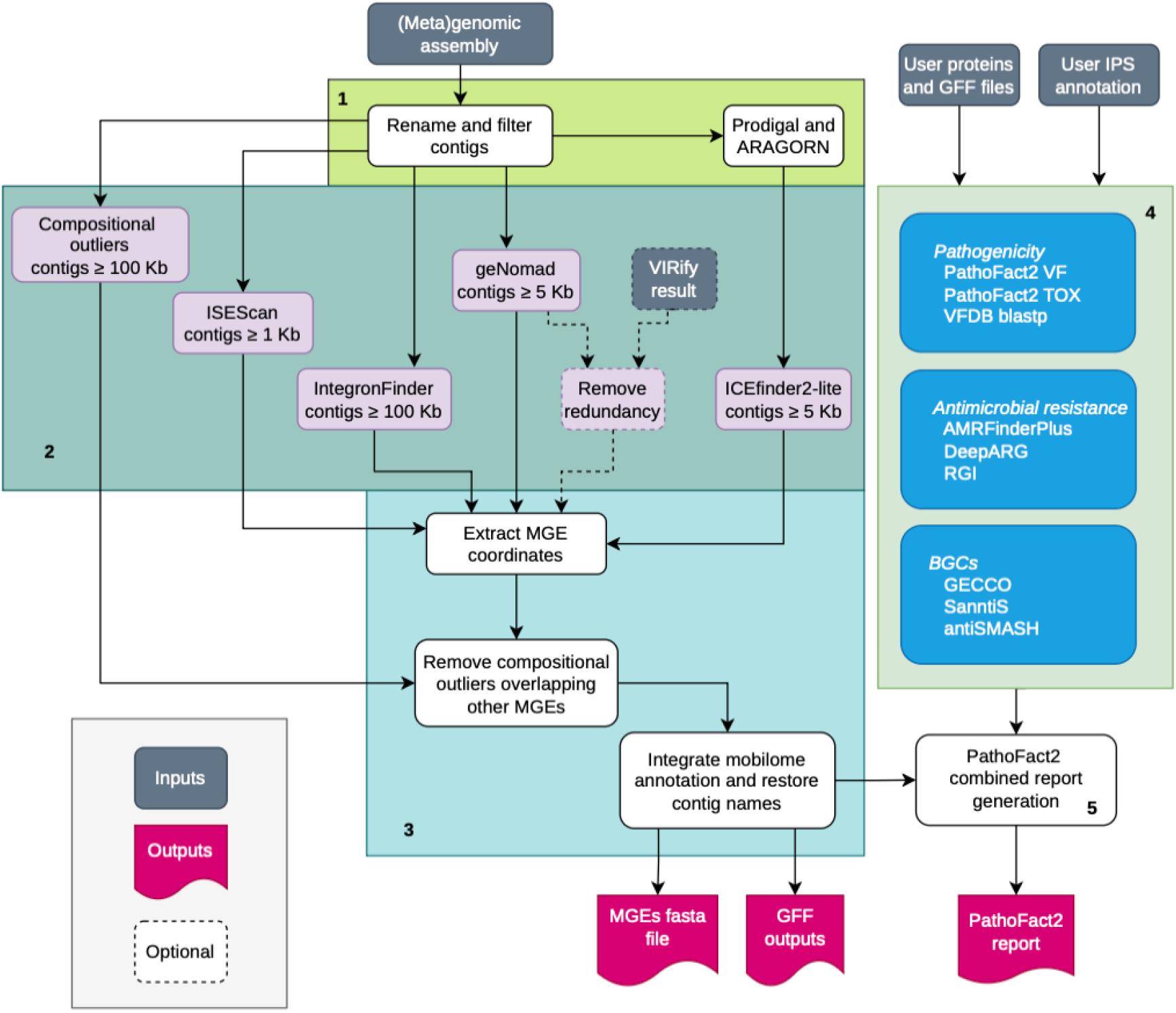
The MAP workflow depicted in 5 blocks of data generation and processing. (1) Preprocessing of the contigs includes renaming, length filtering, and gene/tRNA calling; (2) parallel MGE prediction using multiple tools; (3) integration into a mobilome GFF3 file and MGE FASTA; (4) functional annotation of the predicted proteins to detect antimicrobial resistance genes (ARG), virulence factors and toxins (VF/TOX). This block includes a subworkflow to identify biosynthetic gene clusters (BGC); (5) integration of the annotation modules to report ARG and VF/TOC in the context of BGC and the mobilome. Pipeline inputs are shown in grey, outputs in magenta, and optional inputs or steps are dashed.

Three functional-annotation subworkflows run on the predicted proteins (see Figure 1 section 4). Virulence factors (VF) and toxins (TOX) are identified using the PathoFact2 machine-learning modules (Delgado et al. 2026) and a DIAMOND (Buchfink et al. 2021) search against VFDB (Liu et al. 2022). Antimicrobial resistance genes (ARGs) are called by AMRFinderPlus (Feldgarden et al. 2021), DeepARG (Arango-Argoty et al. 2018) and RGI/CARD (Alcock et al. 2023) and the format is standardised with hAMRonization (Mendes et al. 2024). BGCs are predicted by GECCO (Carroll et al. 2021), SanntiS (Sanchez et al. 2023) and antiSMASH (Blin et al. 2025) and overlapping coordinates are merged across tools. SignalP (Teufel et al. 2022) annotations are extracted from InterProScan results. These annotations are collected in a final output that reports each protein annotated as resistance or virulence in the BGC and MGE context (Figure 1 step 5).

## 3 Use case

### 3.1 Dataset and analysis design

To illustrate MAP utility we applied it to a selection of 421 genomes (a mix of metagenome assembled genomes (MAGs) and isolate genomes including plasmids) from the MGnify soil genome catalogue v1.0 (Gurbich et al. 2023) available at https://ftp.ebi.ac.uk/pub/databases/metagenomics/mgnify_genomes/soil/v1.0/. The pipeline performance stats are available in the Supplementary Table 1. We re-annotated these genomes with MAP v5.0.0, reusing the catalogue’s AMRFinderPlus, SanntiS, GECCO and antiSMASH outputs through the annotation manifest and adding VIRify v4.0.0 predictions. Using the combined report generated by the MAP, we identified 30 genomes (16 MAGs and 14 isolates) carrying a large number of ARG and VF/TOX genes in the mobilome (Figure 2A; Supplementary Table 2). We selected an illustrative set of 43 plasmid contigs of length > 120 kb carrying more than 15 combined ARG and VF/TOX genes to search against public metagenomes with the MGnify-implemented Branchwater v0.4.0 database v2024-11-28 (Irber et al. 2022) to assess their prevalence across different biomes. Only near-complete matches with sequence containment >=0.90 and containment Average Nucleotide Identity (cANI) reported >=0.99 were considered. Detections were collapsed to distinct BioProjects per biome to avoid sampling-effort bias.

**Figure 2.**
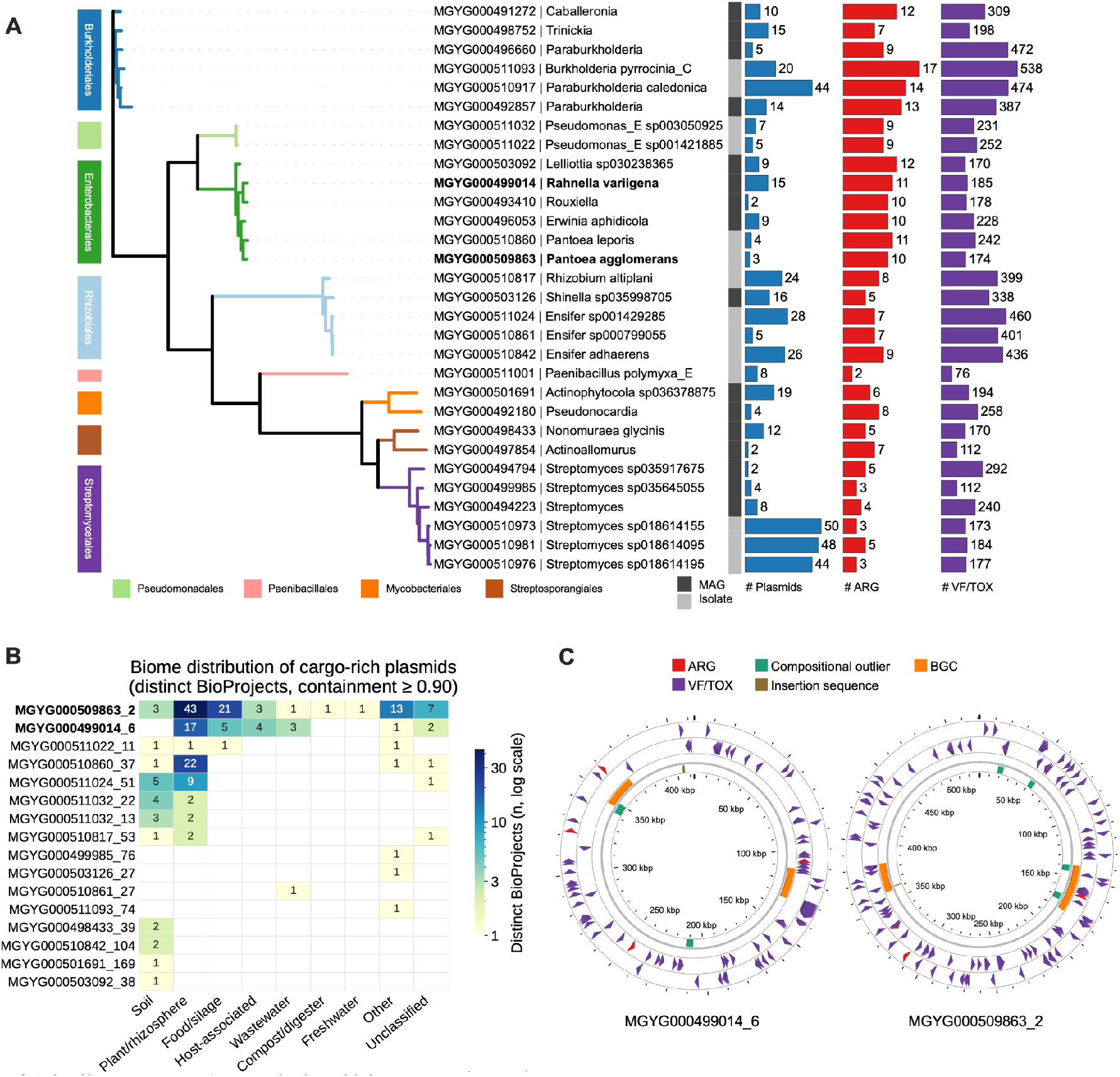
MAP applied to cargo-rich plasmids from the MGnify soil catalogue. (A) A maximum-likelihood phylogeny of the 30 genomes carrying the largest number of ARG/VF/TOX in the mobilome (GTDBtk v2.7.1 (Chaumeil et al. 2022) database release 232; IQ-TREE v2.2.0.3 (Minh et al. 2020) with LG+R10 model) of the 30 plasmid-carrying genomes, coloured by taxonomic order and assembly type (MAG or isolate) with per-genome bars for plasmid count and mobilome ARG and VF genes (this includes toxins); the two hosts examined in detail (*Pantoea agglomerans* and *Rahnella variigena*) are in bold. (B) Biome distribution of the 16 plasmids (containment >=0.90) recovered from public metagenomes, coloured by the number of distinct BioProjects per biome (log scale). (C) Circular CGView (Stothard et al. 2019) maps of the two selected plasmids: BGCs (orange), cargo CDSs (VF/TOX purple, ARG red) and mobile elements (compositional outliers green, insertion sequences brown).

### 3.2 Cross-biome prevalence of cargo-rich plasmids

Sixteen out of the 43 query plasmids from soil metagenomes (6 plasmids from MAGs and 10 plasmids from isolates) were recovered beyond their source BioProject, most often in plant/rhizosphere (51%) and food/silage (14%) metagenomes, with smaller contributions from host-associated (4%) and wastewater (3%) samples (Figure 2B; Supplementary Table 3).

We examined in greater detail the two most widely distributed plasmids, both from Enterobacterales hosts. A 521 kb plasmid from a *Pantoea agglomerans* isolate (MGYG000509863_2) showed near-identical copies (cANI 0.99-1.00) in 82 BioProjects from 9 different biomes across 16 countries, whereas a 413 kb plasmid from a *Rahnella variigena* MAG (MGYG000499014_6) had a narrower, but more biome-diverse footprint that included bovine gut and bat metagenomes. Repeating the search with the complete host genomes gave concordant distributions: every plasmid-positive sample also contained the near-identical host genome and none carried the plasmid alone, so the cross-biome occurrence reflects clonal dissemination of the host strains rather than independent plasmid transfer. A sample of the MAP annotations of the two plasmids depicted in Figure 2C, shows virulence-factor/toxin genes distributed around each replicon (76 and 11 respectively) with a few resistance genes (4 and 3 respectively), and an ARG part of a BGC in each plasmid.

In summary, MAP integrates tools for the prediction of a broad prokaryotic MGE repertoire with functional annotation of its resistance, virulence and biosynthetic gene cluster cargo in a single reproducible and portable pipeline, producing a validated GFF3 file and a combined report that places virulence-related genes in their mobilome and BGCs context. MAP has recently been incorporated into the MGnify suite of analysis pipelines and is used to annotate the mobilome of the MGnify genome catalogues and metagenomic assemblies.

## Data availability

The MAP pipeline and the scripts used to generate the analyses and figures are available at https://github.com/EBI-Metagenomics/mobilome-annotation-pipeline. The MAP execution trace file for the 421 selected genomes is provided in Supplementary Table 1. Genome and plasmid accessions and the Branchwater search results are provided in Supplementary Tables 2 and 3.

